# Neuronal stop-codon readthrough is associated with ribosome pausing and appends lineage-restricted extensions that can alter protein localization in *Drosophila*

**DOI:** 10.64898/2026.07.13.738141

**Authors:** Takashi Makino, Koya Sakuma, Hiroto Anbo, Yilin Zhang, Shilin Lu, Cai Xu, Shintaro Iwasaki, Satoshi Fukuchi, Motonori Ota, Toshiharu Ichinose

## Abstract

Stop-codon readthrough (RT) diversifies proteomes and is particularly prominent in neurons. However, the regulatory logic that specifies neuronal RT and the structural and cellular consequences of the resulting C-terminal protein extensions remain poorly understood. Here we leverage neuron-specific ribosome profiling datasets in *Drosophila* to construct an *in vivo* atlas of 163 neuronal RT transcripts. Sequence-based modeling distinguished RT from non-RT transcripts and highlighted an extended post-stop region enriched for stable predicted RNA structures. Beyond these *cis-*associated features, ribosome-footprint analysis revealed pronounced stop-codon pausing on RT transcripts, accompanied by upstream periodic peaks consistent with ribosome queuing. At the protein level, neuronal RT appended polypeptides enriched in polar residues and intrinsically disordered regions (IDRs). Evolutionary comparisons suggested that most analyzed extensions show shallower phylogenetic conservation than their associated conventional proteins, whereas in two genes a stop-codon appears to have been gained within ancestral coding sequences. Finally, an *in vivo* dual-color reporter showed that RT of the RNA-binding protein Bru3 alters localization from the nucleus to cytoplasmic granules. Together, our results suggest that neuronal RT is associated with structured post-stop RNA regions that reshape termination dynamics and produce lineage-restricted, IDR-rich C-terminal protein extensions with distinct subcellular localization.

**Key points:**

- An atlas of 163 readthrough transcripts in *Drosophila* neurons reveals enhanced stop-codon pausing.
- Readthrough appends IDR-rich, lineage-restricted C-terminal extensions to conserved protein cores.
- Readthrough alters Bru3 localization, enriching its extended form in cytoplasmic granules.

## Introduction

Stop-codon readthrough (RT) occurs when ribosomes bypass a stop codon and continue translation, generating C-terminally extended protein isoforms ^1^. Because RT allows the same transcript to encode both the standard protein and a distinct C-terminally extended variant, it offers an economical route to proteome diversification. Genome-wide approaches, such as ribosome profiling, have revealed widespread stop-codon RT across eukaryotes, from budding yeast ^2^ to *Drosophila* ^3,4^ and mammalian systems ^5–8^. In some systems, RT is often described as being strongly shaped by a short local termination context: UGA is typically the most permissive stop codon, C at the +4 position has often been reported as particularly readthrough-prone, and +4 to +9 nucleotides are suggested to exert the largest effects ^5,7,9^. In contrast, studies focused on individual genes posit that extended downstream elements as far as ∼100 nt from the stop codon can promote RT ^10,11^. The field also differs on how broadly RT is biologically functional: while some argue that most RT is generally nonadaptive and the product can be swiftly degraded ^12–14^, there are multiple lines of evidence that RT is functionally important ^10,15–20^.

The *Drosophila* nervous system provides a particularly informative context to address these issues, where RT appears enriched and biologically relevant. Comparative genomics indicates that many *in silico*-predicted *Drosophila* RT candidates encode conserved protein-coding sequences beyond annotated stop codons across species, implying evolutionary constraint on RT-derived extensions ^21,22^. Independent lines of evidence across multiple genes suggest that RT occurs more frequently in neurons than in other cell types ^11,15,16^. Despite these advances, we still lack a neuron-resolved and transcriptome-wide view of which endogenous mRNAs undergo RT in *Drosophila*, what C-terminal extensions they encode, and which *cis* features distinguish them.

To address these gaps, we combined neuron-specific ribosome profiling datasets with sequence-based modeling and protein-level characterization. Using this framework, we define a neuron-specific atlas of RT, build a sequence-based predictive model, relate RT to distinctive ribosome behavior at stop codons, and characterize the extension proteoforms that can alter protein localization. Our findings thus provide insight into how stop-codon recoding contributes to neuronal proteome diversification.

## Results

### A neuron-specific atlas of stop-codon readthrough in *Drosophila*

To characterize stop-codon RT in *Drosophila* neurons, we performed neuron-specific ribosome profiling ^23–25^: we expressed FLAG-tagged ribosome protein-L3 (*UAS-RpL3::FLAG*) ^26^ by pan-neuronal driver (*nSyb-GAL4*) followed by immunoprecipitation and ribosome footprinting. We observed enrichment of neuronal marker genes compared to the whole-head sample ^24^ (Figure S1), and clear 3-base periodicity consistent with codon-by-codon ribosome movement (Figure 1A). While translation terminated efficiently at the annotated stop codon on most mRNAs (93.25% and 1.91% of the footprints were detected on CDS and 3’ UTR respectively; Figure 1A-B), a subset of mRNAs showed ribosome footprints downstream of the stop codon consistent with stop-codon RT (Figure 1C-F).

**Figure 1.**
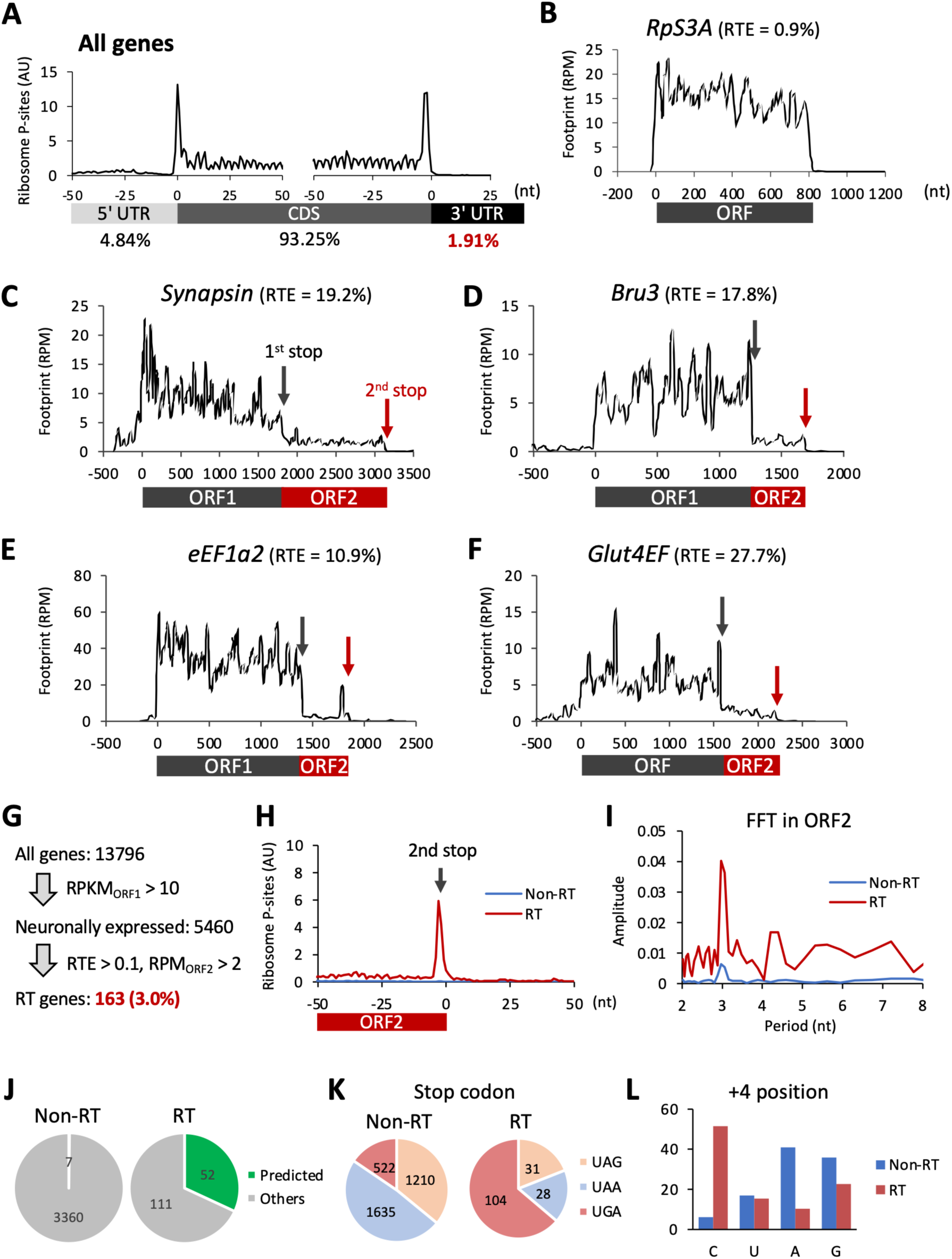
A neuron-specific ribosome-profiling atlas identifies stop-codon readthrough transcripts in *Drosophila* neurons. (A) Meta-genome ribosome distribution (estimated P-sites) around annotated start and stop codons in *Drosophila* neurons. Footprints show clear 3-nt periodicity and are predominantly distributed within coding sequences. The percentages indicate the fraction of ribosome footprints assigned to the 5’ UTR, CDS, and 3’ UTR. (B-F) Representative ribosome-footprint profiles for non-RT- (B) and RT- (C-F) transcripts. The whole footprints on each transcript are plotted. ORF1 indicates the annotated coding sequence and ORF2 the in-frame C-terminal extension between the annotated stop codons (grey arrows) and the next in-frame stop codons (red arrows). RTE, calculated as RPKM_ORF2_/RPKM_ORF1_, is indicated above. (G) Gene classification based on RTE. For all the neuronally expressed genes (RPKM_ORF1_ > 10; 5460 genes) RTE is calculated. 163 genes exhibit RTE > 0.1 and RPM_ORF2_ > 2 and are classified as RT-genes. (H) Ribosome-footprint distribution around the second in-frame stop codon. Estimated ribosome P-sites are aligned to the second stop codon, and footprint densities from −50 to +50 nt are normalized by RPKM_ORF1_ and averaged across transcripts in the non-RT and RT groups. For transcripts with short ORF2 regions (<50 nt), upstream positions that extend into ORF1 are excluded from the average. (I) Three-nucleotide periodicity in ORF2 quantified by Fast Fourier Transform (FFT). FFT amplitudes were calculated from averaged ribosome-footprint densities within the last 100 nt of ORF2 upstream of the second stop codon. For transcripts with ORF2 regions shorter than 100 nt, only the available ORF2 positions were included; positions extending into ORF1 were excluded. (J) Overlap between identified RT genes and previously predicted RT-candidates based on comparative genomics ^21^. (K) Stop codon usage (the annotated stop codon) among RT or non-RT transcripts. (L) Nucleotide frequences at the +4 position (immediate downstream of the annotated stop codon) in RT or non-RT transcripts.

To define neuronal RT genes, we computed readthrough efficiency (RTE), defined as the ratio of ribosome density in the second ORF (i.e., region from right downstream of the first stop codon to the next stop codon) to that in the first ORF (RPKM_ORF2_ /RPKM_ORF1_). Among 5,460 neuronally expressed genes (RPKM_ORF1_ > 10), 163 genes (3.0%) met our thresholds (RTE > 0.1 and RPM_ORF2_ > 2) and were designated as RT genes (Figure 1G) (Supplementary Table 1). Genes with RTE < 0.1 were designated as non-RT controls. On RT transcripts, we observed a prominent footprint peak at the second in-frame stop codon (Figure 1H) and a strong 3-base periodicity in the second ORF (Figure 1I), consistent with in-frame translation of the second ORF. The identified RT genes were significantly enriched among previously predicted *Drosophila* RT candidates: ∼32% of the identified RT genes were predicted by sequence conservation among *Drosophila* species while only ∼0.2% for the non-RT controls (Figure 1J) ^21^. Consistent with prior works ^5,7,9^, UGA stop codon and C at the +4 position occurred more frequently at RT sites than in non-RT controls (Figures 1K and 1L). Altogether, these analyses define a neuron-resolved set of stop-codon RT transcripts in *Drosophila*.

### Sequence modeling reveals an extended post-stop signature associated with neuronal RT

We next asked whether the broader sequence context surrounding the stop codon could distinguish RT from non-RT transcripts. We trained a three-layer 1D convolutional neural network (CNN) (Figures 2A and S2) using sequences from 131 randomly selected RT and 131 non-RT transcripts spanning -100 to +100 nt around the annotated stop codon. We then evaluated the trained model on an independent held-out validation set. Across five independent runs with different random seeds, the model achieved 88.8%±1.4% accuracy, 91.3%±4.0% precision, and 86.1%±4.6% recall (Figure 2A). These results indicate that this 200-nt sequence window contains substantial information that distinguishes RT from non-RT transcripts in our dataset.

**Figure 2.**
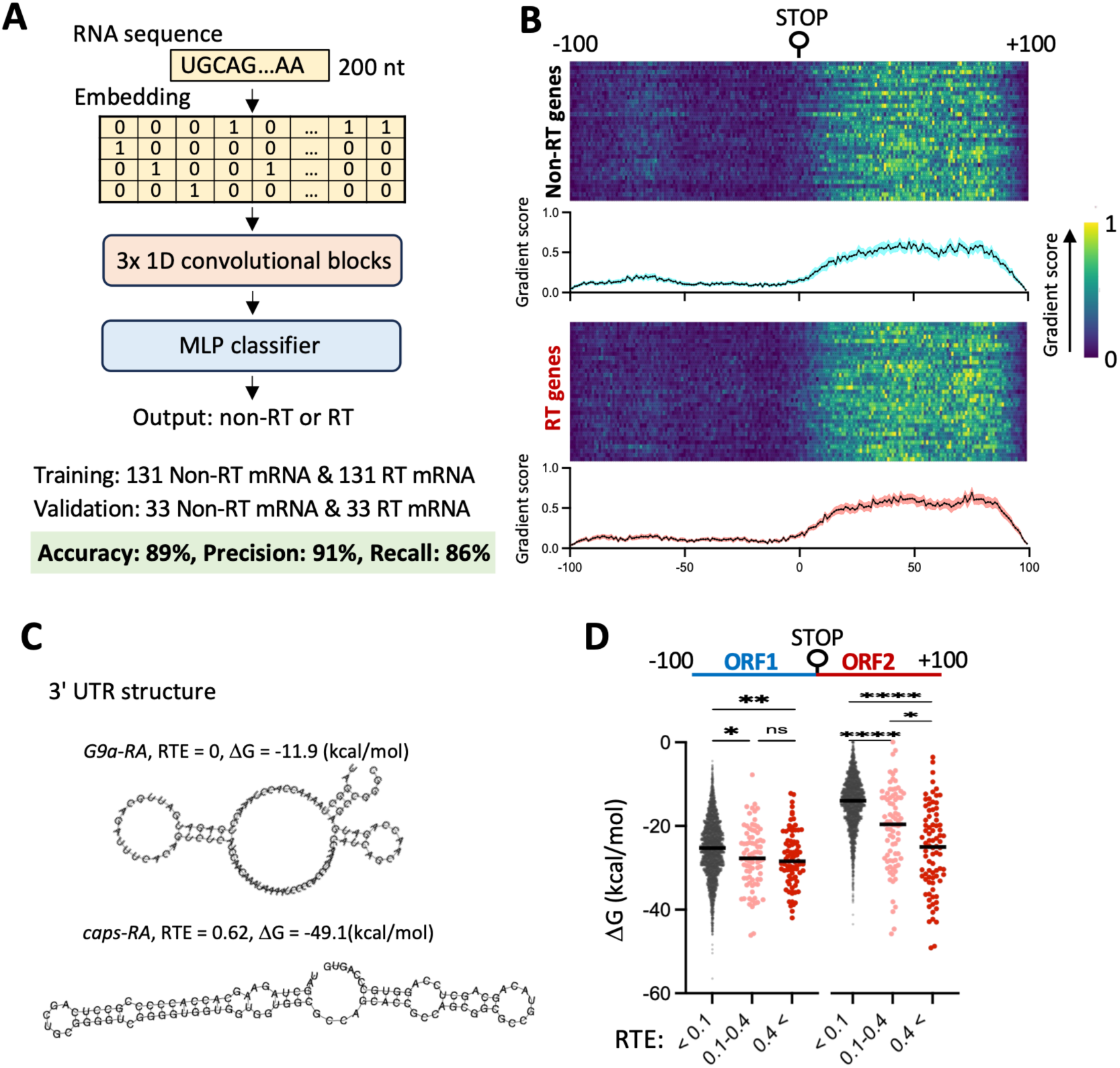
Post-stop sequences and predicted RNA structures distinguish neuronal RT transcripts. (A) Schematic of our 1D convolutional neural network used to classify RT- and non-RT transcripts from nucleotide sequence alone. See also figure S2 and the methods section for more details. Input sequences span 200 nt centered on the annotated stop codon. The model was trained on 131 RT and 131 non-RT transcripts and evaluated on an independent validation set of 33 RT and 33 non-RT transcripts (including the sequences of estimated double-RT). (B) Gradient-based attribution analysis of the trained CNN on the held-out test set. Heatmaps show nucleotide-level gradient scores for non-RT and RT transcripts. Each row indicate each transcript and column the position around the stop codon. Line plots at the bottom show the average ± standard error across positions. (C) Representative RNA secondary-structure predictions (RNAfold) for 100-nt sequences in the 3’ UTR of a non-RT transcript, *G9a-RA*, and an RT transcript, *caps-RA*. Minimum free-energy values are shown. (D) Predicted minimum free energy (RNAfold) for sequences upstream and downstream of the annotated stop codon, grouped by RTE. RT transcripts show lower predicted free energy in the post-stop region, consistent with stronger RNA secondary structure. Dunn’s multiple comparison test is used and center lines indicate the median. ns, not significant; \**P* < 0.05; \*\**P* < 0.01; \*\*\*\**P* < 0.0001.

To identify which nucleotide positions contributed to the CNN classification, we performed gradient-based attribution analysis. In this analysis, the contribution of each input nucleotide is estimated from how sensitively the CNN output, here the predicted probability of RT, depends on the nucleotide encoded at that position ^27,28^. This analysis demonstrated that nucleotides downstream of the stop codon, extending to ∼100 nt, dominated the model’s predictions in both non-RT and RT prediction (Figures 2B and S3), highlighting the relevance of this region for RT. Because several gene-level studies have shown that stem-loop structures downstream of stop codons can promote RT in *Drosophila* ^10,11^, we next predicted RNA secondary structure using RNAfold ^29,30^. RT transcripts, especially those with high RTE (> 0.4), exhibited markedly higher structural stability (lower minimum free energy) downstream of the stop codons than non-RT controls, whereas upstream differences were modest (Figure 2C-D). Together, the sequence-based classifier and structure predictions implicate secondary structure downstream of the stop codon as a prominent feature associated with neuronal RT in *Drosophila*.

### Neuronal RT sites show stop-codon pausing and upstream ribosome queuing

Because stem-loop structures can impede elongating ribosomes and thereby influence ribosome kinetics ^31^, we hypothesized that ribosome dynamics differ between RT and non-RT transcripts. Although ribosome footprints generally accumulated at both start and stop codons (Figure 1A), stop-codon-associated pausing was markedly enhanced on RT transcripts and was strongest in transcripts with high readthrough efficiency (RTE > 0.4) (Figures 3A and 3C). In contrast, the start-codon peak did not differ significantly between RT and non-RT transcripts (Figures 3A and 3B). Consistent with enrichment of UGA among RT sites (Figure 1I), pausing was strongest at UGA and moderate at UAA (Figure 3D-F). Among non-RT transcripts, a subtle but statistically significant increase of ribosome pausing at UGA was also observed (Figure S4).

**Figure 3.**
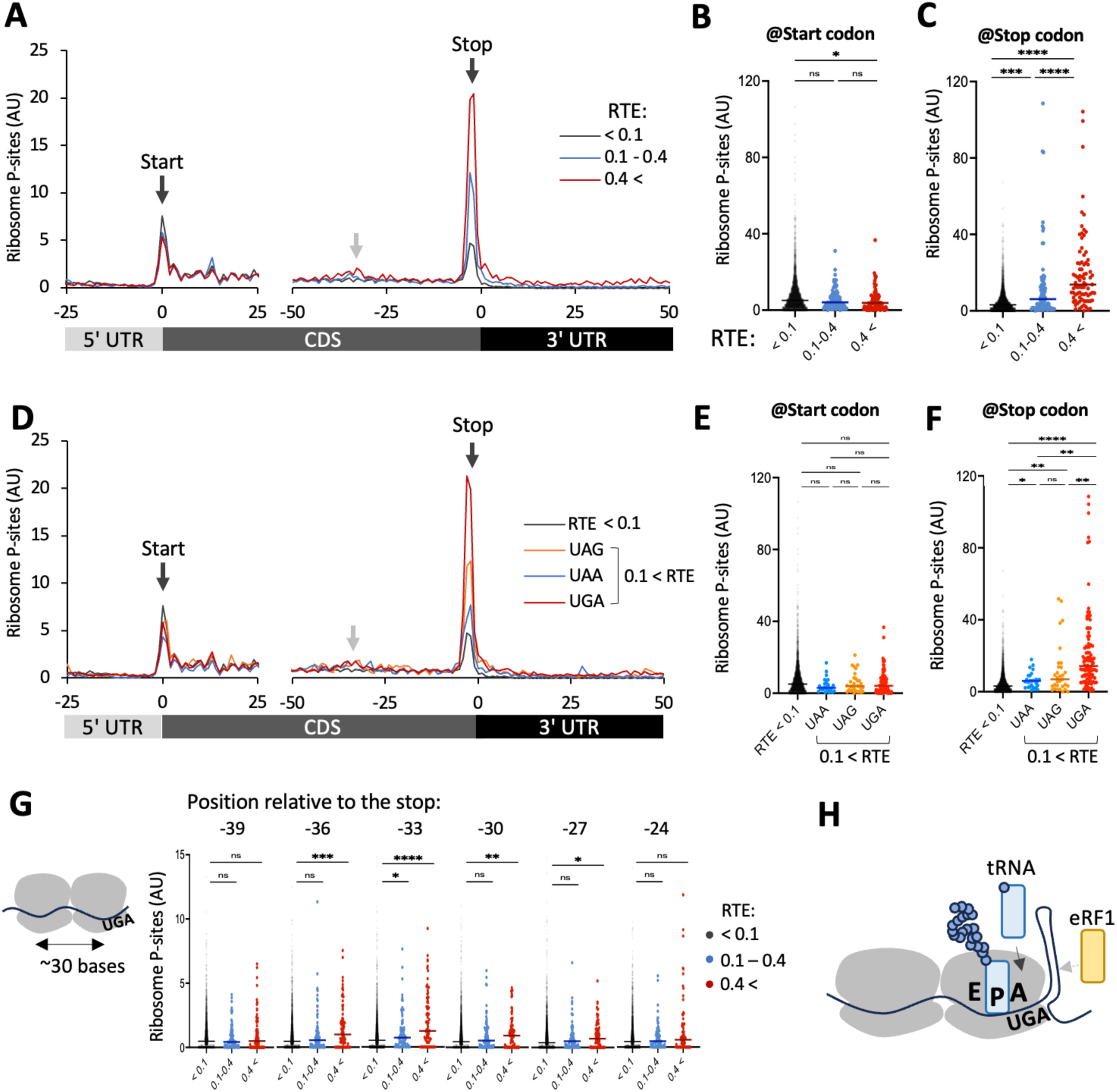
Readthrough transcripts show stop-codon pausing and upstream ribosome queuing. (A) Metagene profiles of ribosome P-sites around start and stop codons, stratified by RTE. (B, C) Quantification of ribosome P-site density at the start codon (B) and at the position 3 nt upstream of the annotated stop codon (C), corresponding to a ribosome with the stop codon positioned in the A-site. (D) Metagene profiles stratified by RTE and stop-codon identity. (E, F) Quantification of ribosome P-site density at the start codon (E) and at the stop-codon −3 position (F), grouped by stop-codon identity and RTE. (G) Quantification of ribosome P-site density at upstream positions spaced approximately one ribosome footprint apart from the stop codon. RT transcripts showed elevated ribosome density near −33 nt and neighboring positions, consistent with upstream ribosome queuing. (H) Model illustrating ribosome pausing at the stop codon and competition between release-factor-mediated termination and near-cognate tRNA-mediated readthrough. Dunn’s multiple comparison test is used, and center lines indicate the median. ns, not significant; \**P* < 0.05; \*\**P* < 0.01; \*\*\**P* < 0.001; \*\*\*\**P* < 0.0001.

In addition, we noticed that RT transcripts displayed periodic upstream peaks at approximately -33 nt and -66 nt relative to the stop codon (grey arrows in Figures 3A and 3D; see also Figure S5). Ribosome footprints were significantly higher at these positions on RT-transcripts (Figures 3G and S5). Given that a ribosome footprint spans around 30 nucleotides ^32^, these peaks are consistent with ribosome queuing behind a stalled ribosome at the stop codon. Altogether, these results indicate that ribosomes undergo strong pausing at RT sites, particularly at RT-prone UGA stop codons (Figure 3H).

### C-terminal extensions are enriched for intrinsically disordered regions

We next investigated the properties of proteins encoded by RT transcripts. Gene Ontology (GO) enrichment analysis identified only one significantly enriched term, “positive regulation of transcription” (Figure S6), whereas UniProtKB sequence feature analysis revealed significant enrichment for several terms, including “polar residues” and “disordered regions” (Figure 4A). Because these annotated sequences do not necessarily include the RT-derived ORF2 extensions, we directly examined the amino-acid composition in ORF1- and ORF2-encoded sequences. Relative to the non-RT controls, polar and electrically-neutral residues such as Gln and His were strongly enriched in the ORF2 of RT transcripts (108.9% and 57.5% increase for Gln and His, respectively), and also slightly in the ORF1 (22.6% and 22.0% for Gln and His, respectively) (Figure 4B).

**Figure 4.**
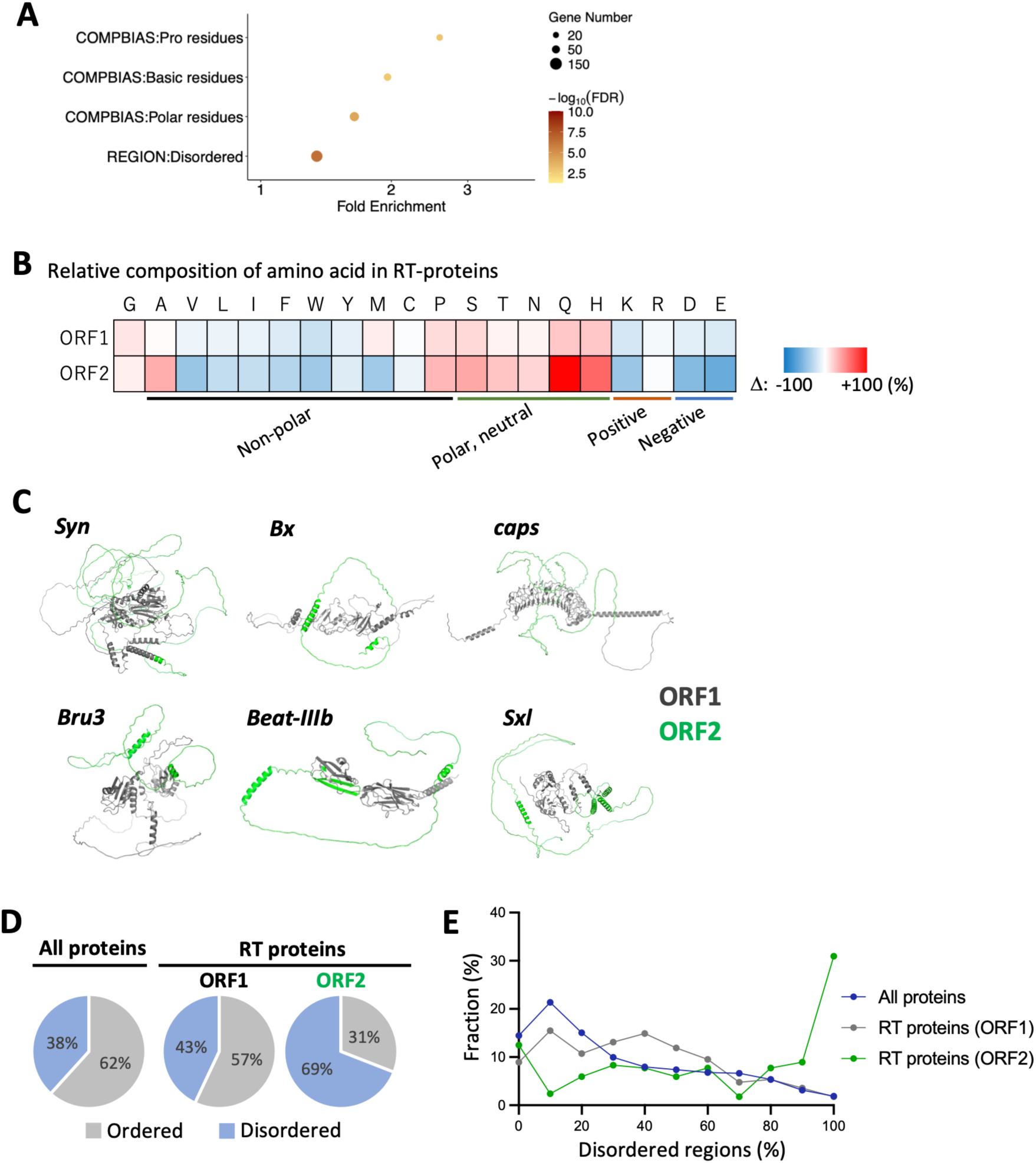
Stop-codon readthrough appends intrinsically disordered C-terminal extensions. (A) Enrichment analysis of UniProtKB sequence features among RT proteins. RT proteins are enriched for features associated with disordered regions and compositionally biased regions, including polar, basic, and proline-rich residues. Dot size indicates gene number, and color indicates −log 10 (FDR). (B) Relative amino-acid composition of ORF1 and ORF2 regions in RT proteins. Enrichment of amino acid compared to ORF1 in non-RT controls is shown with a color code. (C) Representative AlphaFold2-predicted structures of RT proteins. ORF1 regions are shown in grey and readthrough-derived ORF2 extensions in green. Many ORF2 extensions adopt extended conformations consistent with IDRs. (D) Fraction of ordered and disordered regions in all proteins, ORF1 regions of RT proteins, and ORF2 extensions of RT proteins, predicted by NeProc ^35^. (E) Distribution of predicted disordered-region content in all proteins (blue), RT-protein ORF1 regions (grey), and RT-protein ORF2 extensions (green).

To examine structural characteristics, we predicted RT-protein structures with AlphaFold2 or ColabFold ^33,34^. We found that many RT-encoded tails exhibited extended conformations consistent with intrinsically disordered regions (IDR) (Figure 4C) (supplementary files for all RT proteins). To quantify predicted disorder more directly, we applied a computational model we previously developed (NeProc) ^35^. Compared with proteome-wide baseline (IDR: 38%), we observed modestly elevated IDRs in ORF1 segments of RT proteins (43%) and a pronounced increase in ORF2 extensions (69%) (Figure 4D). Notably, our model predicted that 39.9% of C-terminal extensions (ORF2) have more than 90% disordered residues, compared with 5.0% for all proteins and 5.4% for ORF1 segments in RT transcripts (Figure 4E). Taken together, these data indicate that neuronal RT appends C-terminal tails enriched with IDRs.

### Neuronal RT extensions show shallow phylogenetic conservation

Several studies reported elevated intrinsic disorder in proteins encoded by evolutionarily young genes ^36–38^. The high predicted disorder content of ORF2 extensions therefore prompted us to investigate their evolutionary histories relative to their associated ORF1-encoded protein cores. We focused on 87 RT genes with ORF2 extensions longer than 50 amino acids and examined ORF1 and ORF2 conservation across 13 representative arthropod species. We combined protein- and genome-level sequence similarity searches, including translation of downstream genomic regions where protein annotations did not cover the putative extension (Methods).

ORF1 sequences generally showed deep conservation (Figure 5A): 84 of the 87 genes had detectable homologous sequences at the broadest arthropod level examined, whereas the remaining three were assigned to *Cyclorrhapha*, *Diptera*, or *Hexapoda* (Figures 5B and 5C). In contrast, ORF2 conservation was predominantly restricted to shallower clades. For 82 of the 87 genes, ORF2 similarity was detected only within *Diptera*, including 48 for which detection was restricted to *Drosophilnae* or shallower clades. Only two ORF2 sequence yielded a significant hit at the broadest *Arthropoda* level (Figures 5B and 5C). Thus ORF2 extensions generally exhibited a more restricted phylogenetic distribution of sequence similarity than their cognate ORF1 regions.

**Figure 5.**
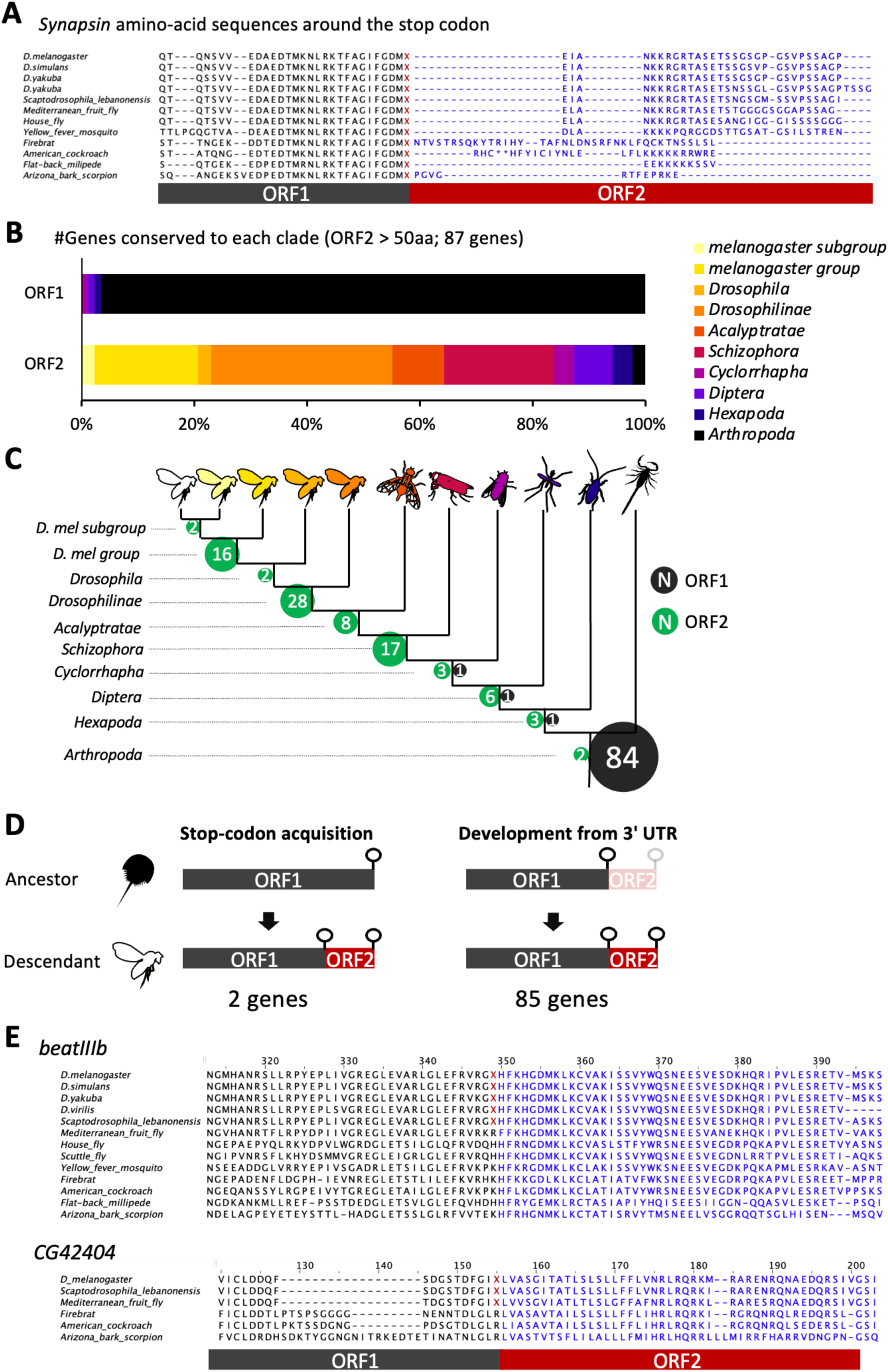
Evolutionary conservation and inferred origins of neuronal RT extensions. (A) Cross-species sequence alignments of Synapsin, showing stronger conservation of ORF1 than of ORF2. (B) Distribution of conservation depths for ORF1 and ORF2. Homologous sequences of ORF1 and ORF2 were examined across 13 arthropod species, and were assigned to the deepest phylogenetic clade with detectable sequence similarity (Methods). Stacked bars show the percentage of genes assigned to each clade. 87 neuronal RT genes with ORF2 extensions longer than 50 amino acids are analyzed. (C) Conservation categories from (B) mapped onto a schematic phylogeny. Numbers indicate the genes assigned to each clade for ORF1 (black) and ORF2 (green). (D) Two proposed evolutionary routes to RT-derived extensions. Left, acquisition of an internal stop codon within an ancestral continuous CDS makes translation of a pre-existing C-terminal region dependent on RT. Right, an ancestral 3′ UTR gradually evolves into a translated C-terminal extension. Sequence comparisons supported the stop-codon-gain model for two genes. The remaining 85 genes lacked a comparable phylogenetic pattern of a continuous CDS in distant species. (E) Sequence alignments spanning the ORF1–ORF2 boundary in *beat-IIIb* and *CG42404*, which are consistent with the acquisition of an internal stop codon. Blue letters indicate residues corresponding to the *D. melanogaster* ORF2 region, and red X characters mark positions as stop codons.

We next considered two possible evolutionary routes to the present-day RT architecture (Figure 5D). In one scenario, an internal stop codon is gained within an ancestral coding sequence, making translation of a pre-existing C-terminal region dependent on RT. In the other, part of the ancestral 3’ UTR became translated through RT, generating a new C-terminal extension. To distinguish these possibilities, we examined whether sequences corresponding to the *D. melanogaster* ORF1 and ORF2 regions formed a continuous coding sequence in other species or were separated by a stop codon.

This analysis identified two genes, *beat-IIIb* and *CG42404*, for which sequences corresponding to ORF1 and ORF2 formed a continuous coding sequence in distant species (Figure 5E). This result supports the gain of an internal stop codon within an ancestral CDS. For the remaining 85 genes, we found no comparable pattern of a continuous CDS in distant species, consistent with the second scenario. Together, these analyses suggest that many neuronal RT extensions are lineage-restricted and 3’ UTR-derived additions to deeply conserved protein cores, with two exceptions in which an internal stop-codon appears to have ben gained within an anscentral CDS.

### Readthrough alters the *in vivo* subcellular localization of Bru3

Because subcellular localization signals are often encoded within IDRs ^39,40^, we next tested whether RT alters protein localization in *Drosophila* neurons. We generated dual-color reporters for five genes with relatively high RTE and known molecular functions: *eEF1a2* (RTE = 0.109), *Bruno-3* (*Bru3*) (0.178), *Synapsin* (0.192), *Glut4EF* (0.277), and *CASK* (0.461). In these constructs, tagRFP was fused to the N-terminus of the cDNA to label all isofoms, whereas Venus was fused to the C-terminus of ORF2 to specifically report the readthrough products (Figure 5A). When expressed using a pan-neuronal driver (*nSyb-GAL4*), we detected robust readthrough Venus signal for *Bru3*, *Synapsin*, *Glut4EF*, and *CASK*, but not for *eEF1a2* (Figure S7). *Synapsin*, *Glut4EF*, and *CASK* showed broadly similar patterns between tagRFP and Venus signals (Figure S7). For the *Bru3* reporter (Figure 5B), both tagRFP and Venus signals were largely restricted to cell bodies (Figure 5C), but their subcellular localization exhibited clear difference: tagRFP was detected in both nucleus and cytoplasm whereas Venus-marked RT products accumulated almost exclusively in cytoplasmic granules (Figure 5D). These observations indicate that RT alters the subcellular localization of Bru3 in neurons *in vivo*.

## Discussion

In this study we generated a neuron-resolved atlas of stop-codon readthrough in *Drosophila*, identifying 163 readthrough genes (Figure 1). A central finding is that RT transcripts display pronounced ribosome accumulation at stop codons and increased predicted stability of downstream RNA structures (Figures 2-3). Furthermore, we found that ORF2 generally encodes polypeptides enriched for IDRs and shows shallower phylogenetic conservation than ORF1 (Figures 4-5). Genetic reporter revealed that RT can alter subcellular localization of Bru3 in mature neurons (Figure 6). Together, these observations support a model in which downstream mRNA architecture contributes to a termination bottleneck that may facilitate near-cognate decoding, while RT generates IDR-rich C-terminal additions that diversify neuronal proteoforms.

**Figure 6.**
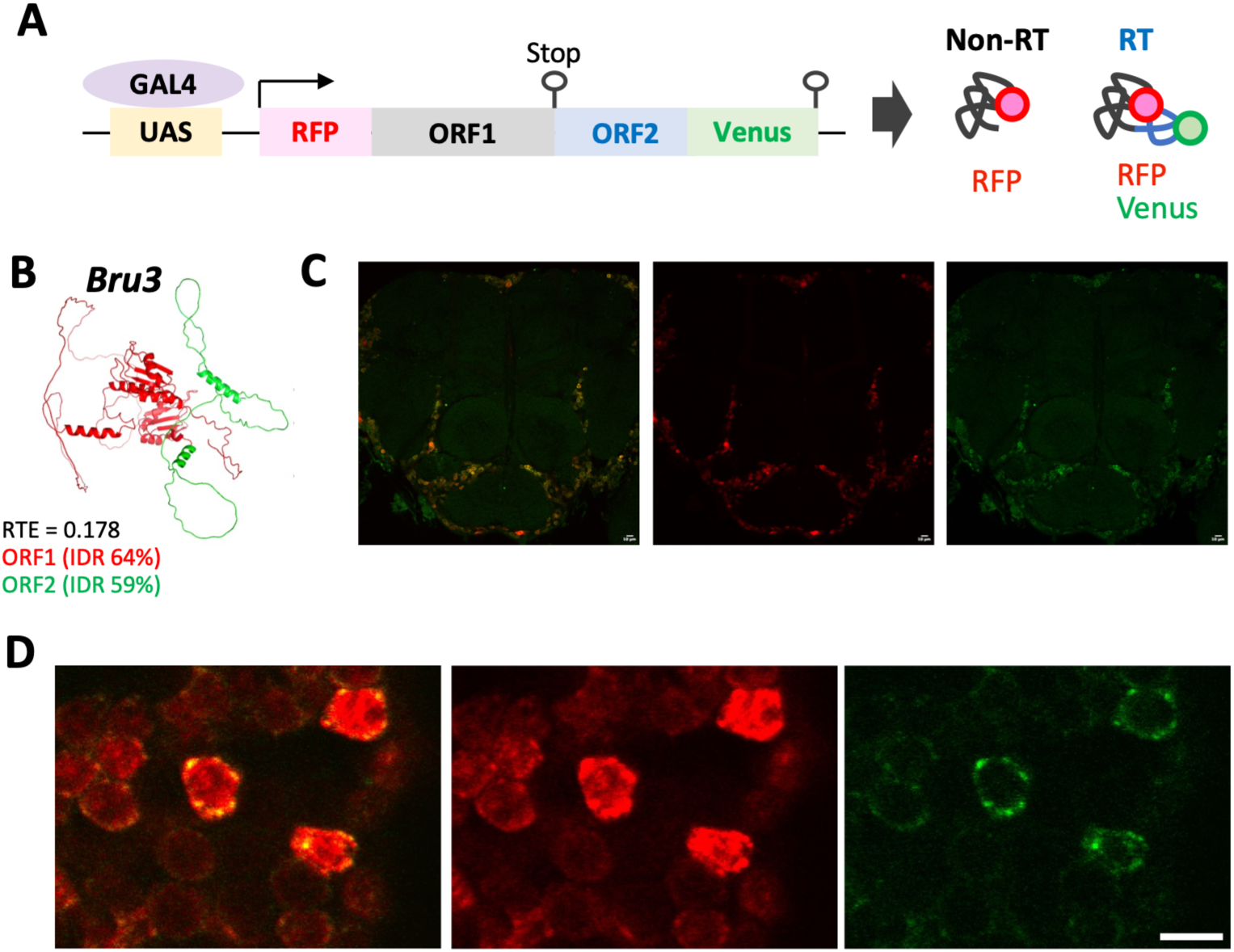
Readthrough redirects Bru3 into cytoplasmic granules in neurons. (A) Schematic of the dual-color readthrough reporter. *tagRFP* is fused to the N-terminus of the coding sequence to label both canonical and readthrough products, whereas *Venus* is fused to the C-terminus of ORF2 to specifically label readthrough products. (B) Predicted structure of the Bru3-RE readthrough product. Polypeptides encoded in ORF1 and ORF2 are shown in red and green, respectively. (C) Representative whole-brain confocal images of the *Bru3* dual-color RT reporter driven by *nSyb-GAL4*. Merged, RFP, and Venus channels are shown. Scale bar, 10 µm. (D) Representative sliced confocal images of the *Bru3* dual-color RT reporter driven by *nSyb-GAL4*. Kenyon cells, the mushroom body intrinsic neurons, are shown. Scale bar, 5 µm.

### Stop-proximal and extended downstream features jointly characterize neuronal RT

Our analyses identify RT-associated *cis* features at multiple spatial scales. At the immediate termination context, neuronal RT sites were enriched for UGA stop codons and C at the +4 position, consistent with established determinants of readthrough in other systems (Figure 1) ^2,5,7,9^. Beyond this local context, CNN attribution scores remained elevated across an approximately 100-nt region downstream of the stop codon, and RT transcripts showed lower predicted minimum free energies in this region (Figure 2). Non-RT transcripts harbor much weaker secondary structures compared to RT (Figure 2D), raising a possibility that there is a constraint to avoid such structures to ensure proper termination. This is consistent with previous gene-level studies in *Drosophila* showing that long downstream cassettes, up to ∼100 nt, play a crucial role in RT ^10,11,41^. Together, these observations suggest that stop-proximal and distal *cis* elements act combinatorially to shape ribosome dynamics at termination sites.

### Ribosome pausing may create a kinetic window for readthrough

Our metagene analyses reveal that RT transcripts exhibit a markedly elevated ribosome footprint peak at the annotated stop codon and two periodic upstream peaks consistent with ribosomes queuing behind a slow leading ribosome. It mirrors collision-aware profiling studies in other systems that identify stop codons as major sites of ribosome collisions ^42–44^, but further elaborates that it is much more prominent on RT stop codons in neurons.

In the standard model of termination, the release factor eRF1 competes with near-cognate tRNAs for the stop codon in the ribosomal A site ^45^. Our observation of extended pausing suggests that on RT transcripts, termination is slowed, thereby widening the time window available for a near-cognate tRNA to successfully decode the stop codon. This mechanistic model aligns with a previous study showing that experimentally reducing termination efficiency promotes readthrough ^2^. The known *trans-*factor for RT, eRF-1H ^16^, may facilitate this process. Altogether, our data suggest that neuronal RT sites are associated with a termination bottleneck that increases the probability of near-cognate tRNA incorporation.

### Distinct evolutionary routes to RT-mediated proteoform diversification

The two evolutionary routes considered here (Figure 5D) have distinct implications for proteoform diversification. For *beat-IIIb* and *CG42404*, our comparisons support the gain of an internal stop codon within an ancestral continuous CDS (Figure 5E). In this scenario, the potential advantage of RT lies not in generating a new C-terminal sequence, but in enabling the production of two proteoforms from a single mRNA. Termination at the newly acquired stop codon produces a shorter protein, whereas RT preserves production of a longer form containing the ancestral C-terminal region. Regulating the balance between these forms could allow their functions or localization to diverge without requiring separate transcripts.

For the remaining genes examined, we found no phylogenetic pattern of stop-codon gain, which is consistent with the alternative model in which ORF2 develops from 3′ UTR (Figure 5D). Under this model, initially low-level RT could expose downstream sequences to selection as C-terminal peptides while maintaining production of the canonical and unextended protein. Extensions that confer advantageous interactions or localization could subsequently be retained. This scenario conceptually parallels the proto-gene model of *de novo* gene birth, in which translation provides opportunities for selection on non-genic sequences ^46^. The ORF2-encoded extensions in our dataset show high predicted disorder content (Figure 4), consistent with previous findings in yeast ^47^. This enrichment also parallels the prevalence of disorder at protein termini ^48^ and in segments affected by alternative splicing ^49^. Such IDRs may facilitate the emergence of short interaction or localization motifs without requiring the evolution of a new folded domain ^50–52^. RT may thus provide a route to the gradual acquisition of functional sequences while initially limiting extended proteoforms to a small fraction of the protein products.

*Bru3* may illustrate such functional specialization. Our dual-color RT reporter experiment suggest that RT alters its localization from nuclear to the cytoplasmic granules (Figure 6). *Bruno* family proteins have been implicated in both pre-mRNA splicing and translational repression ^53–57^. Our findings thus raise the possibility that the shorter form contributes to nuclear splicing whereas the extended form participates in cytoplasmic translational repression. RT has been reported to increase under starvation and oxidative stress ^16,58^. Such regulation could rapidly alter the relative production of short and extended forms from existing mRNAs. Although these functional and regulatory possibilities remain to be tested, our results suggest that an IDR-rich RT extension may contribute to functional diversification of proteins.

## Acknowledgement

We thank Dr. Hiromu Tanimoto (Tohoku University) for his generous supports including providing the experimental space and intellectual inputs, Ms. Mai Kanno (Tohoku University), Dr. Yuichi Shichino (University of Tsukuba) and Ms. Mari Mito (RIKEN) for technical assistance to obtain the original ribosome profiling datasets ^23^, and Dr. Shu Kondo (Tokyo University of Science) for technical advice to create the transgenic RT reporters. We also thank the Bloomington *Drosophila* Stock Center (BDSC) and the *Drosophila* Genomics Resource Center (DGRC) for providing transgenic fly strains and plasmids containing cDNA. Confocal imaging was performed in the FRIS CoRE, a shared research environment in FRIS, Tohoku University. Computation was supported by the HOKUSAI SailingShip supercomputer facility at RIKEN. This work was supported by JSPS/MEXT (21K06369, 21H05713, 25H02436, and 25K22463 to T. I., 20H05932 to S. F. and M. O.), JST FOREST (to T. I.), the Uehara Memorial Foundation Grant (to T. I.), the Kishimoto-foundation (to T. I.), the Lotte Foundation (to T. I.), and RIKEN TRIP initiative “TRIP-AGIS” (to S. I.).

## Declaration of generative AI and AI-assisted technologies in the writing process

During the preparation of this work, the authors used generative AI in order to refine the description. After using it, the authors reviewed and edited the content as needed and take full responsibility for the content of the publication.

## Data, code, and materials availability

The neuron-specific Ribo-seq dataset was deposited at NCBI Bioproject (Submission-ID: SUB16575947, BioProject-ID: PRJNA1548294).

The source code and pretrained model weights (Figure 2) were deposited to GitHub (https://github.com/YilinZhang107/neuronal-readthrough-cnn) and archived on Figshare (https://doi.org/10.6084/m9.figshare.34729635).

Predicted protein structures (Figure 4) were deposited to Zenodo (10.5281/zenodo.23527523).

## Author contributions

Conceptualization: TM, SL, MO, TI.

Methodology: TM, KS, HA, YZ, CX, SI, SF, MO, TI.

Investigation: TM, KS, HA, YZ, TI.

Visualization: KS, YZ, TI.

Funding acquisition: SI, SF, MO, TI.

Project administration: TI.

Supervision: SF, MO, TI.

Writing – original draft: TI.

Writing – review & editing: TM, KS, HA, YZ, SL, CX, SI, SF, MO, TI.

## Methods

### Calculation of readthrough rate and isolation of RT genes

To calculate readthrough rate (RTE), we performed neuron-specific Ribo-seq ^24^. Briefly, a flag-tagged ribosomal protein L3 was expressed in neuronal cells by crossing females of *w;;UAS–RpL3–FLAG* ^26^ (Bloomington *Drosophila* Stock Center (BDSC) #77132) and males of *w;;GMR57C10-GAL4* (*nSyb-GAL4*) (BDSC #39171). Heads of the F1 flies were isolated after flash freezing with liquid nitrogen ^59^, homogenized, immunoprecipitated, and digested with RNase I ^24^. The size-selected ribosome footprints were converted into sequencing libraries and were analyzed with HiSeqX_Ten (Illumina).

We pooled 8 libraries, in total 36,265,580 mapped unique reads, all from *w;;UAS–RpL3–FLAG/GMR57C10-GAL4* flies. Fragments ranging from 21-36 nt were analyzed, and the position of the P-site was estimated as 12 nt downstream of the 5’ end of the reads. RPKM was calculated for the main ORF (ORF1) and the second ORF (ORF2), which was defined as the region between the first and the second in-frame stop codons.

Neuronally expressed genes were first filtered with the threshold of RPKM_ORF1_ > 10. When there are multiple splice isoforms the representative isoform, one exhibiting the biggest RPKM_ORF1_, was selected. Among them, readthrough rate (RTE) = RPKM_ORF2_/RPKM_ORF1_ was calculated and those showing RTE < 0.1 were classified as non-RT mRNAs. Transcripts with RTE > 0.1 and RPM_ORF2_ < 2 were filtered out to take only the reliable RT mRNAs. Transcripts with RTE > 0.1 and RPM_ORF2_ > 2 were classified as RT mRNAs. A few mRNAs whose ORF2 overlaps with neighboring genes and the corresponding ribosome footprints cannot be distinguished were discarded.

These RT mRNAs were further classified according to the identity of stop codons or RTE (more or less than 0.4; Figure 1-3). RT-transcripts are listed in Supplementary Table 1.

### FFT analysis of ribosome periodicity in ORF2

To quantify 3-nt periodicity in the readthrough extension, estimated ribosome P-site densities were analyzed by Fast Fourier Transform (FFT). For each transcript, we extracted footprint counts within ORF2, using up to the last 100 nt upstream of the second in-frame stop codon. Positions extending upstream beyond ORF2 into ORF1 were excluded for transcripts with short ORF2 regions. P-site densities were normalized by RPKM_ORF1_, averaged across transcripts within each group, and then subjected to FFT. FFT amplitudes were plotted as a function of period length.

### Training and examination of the convolutional neural network

To evaluate whether local RNA sequence features can predict stop-codon readthrough, we constructed a binary classifier using 200-nt sequences centered on the annotated stop codon. The dataset contained 166 RT-positive (all the RT genes and additional 3 sequences exhibiting a sign of double readthrough) and 662 RT-negative sequences (genes showing RTE = 0). After excluding positive sequences shorter than 200 nt, 164 positive sequences remained. To obtain a balanced dataset, 164 negative sequences were randomly selected. The resulting 328 sequences were divided by stratified random sampling into training and validation sets at an 8:2 ratio, yielding 262 training samples and 66 validation samples.

Each RNA sequence was one-hot encoded as a 200 × 4 matrix, with each nucleotide represented by a four-dimensional binary vector. We trained a three-layer one-dimensional convolutional neural network (1D-CNN) followed by a fully connected classifier to distinguish RT-positive from RT-negative sequences (Figure S2) ^60,61^. Following convolutional feature extraction, the output h(3) is flattened into a vector *u* ∈ ℝ*^d^*, where *d* is the flattened dimension. The classification head consists of a fully connected hidden layer with 256 units:

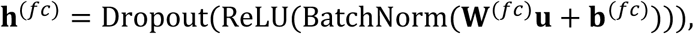

where W(fc) ∈ ℝ^256×*d*^ with dropout rate 0.5. The final output layer produces the classification logit:

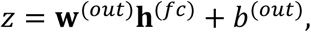

where w(out) ∈ ℝ^256^ and *b*(out) ∈ ℝ. The probability of the positive class is obtained via sigmoid activation: The final fully connected layer contained 256 hidden units followed by a sigmoid output producing the predicted probability *p_i_* of readthrough.

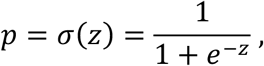

A sequence was classified as positive when *p_i_* > 0.5.

Models were trained by minimizing binary cross-entropy loss,

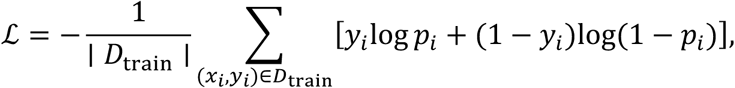

where *y_i_* ∈ {0,1} denotes the class label and *p_i_* = *f_θ_*(*x_i_*) is the model output. Parameters were optimized using the Adam optimizer ^62^ with an initial learning rate of 1 × 10^−4^. A ReduceLROnPlateau scheduler decreased the learning rate by a factor of 0.2 after 7 epochs without improvement in validation area under the receiver operating characteristic curve (AUC-ROC), with a minimum learning rate of 1 × 10^−6^. Training was run for up to 150 epochs with a batch size of 32, and early stopping with a patience of 20 epochs was used to prevent overfitting. The model checkpoint with the best validation AUC-ROC was retained for downstream analysis. To account for variability due to neural network initialization and stochastic training, performance in Figure 2A was reported as the mean ± standard deviation across five independent runs with different random seeds using the same training and validation split. Models were implemented in PyTorch 2.7 and trained on an NVIDIA GeForce RTX 4090 GPU. Model performance was evaluated using accuracy, precision, and recall.

To identify sequence positions that contributed most strongly to CNN predictions, we performed attribution analysis using input gradients ^27,28^. The importance of position j in sequence *X_i_* was calculated as the sum of the absolute gradients across nucleotide channels,

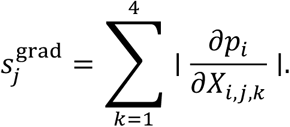

These scores were visualized across training (Figure S3) and test samples (Figure 2B).

### Calculation of RNA secondary structures with RNAfold

RNA secondary structures and the minimum free energies (MFEs; kcal/mol) (Figure 2C-D) were predicted using ViennaRNA package v2.7.0 with default parameters and not structural constraints ^30^. Genes were classified according to RTE and 100 nucleotides before or after the annotated stop codons were used (Figure 2D). For each transcript, we used the same representative isoform as in the Ribo-seq analysis. To compare local RNA-structure stability around annotated stop codons, sequences immediately upstream and downstream of the annotated stop codon were analyzed separately. Specifically, the 100 nt upstream of the stop codon and the 100 nt downstream, including the stop codon itself, were extracted and folded independently (Figure 2D). Transcripts lacking the full 100-nt sequence in either window were excluded from the corresponding analysis. Genes were grouped according to RTE, and the resulting MFEs were compared between groups.

### Prediction of protein structures with AlphaFold2

Protein structure with or without polypeptides derived from ORF2 was predicted using original AlphaFold2 ^33^ or ColabFold ^34^ (Figure 4C). For calculation of the RT polypeptides, stop codon was replaced with Trptophan (encoded by UGG). AlphaFold2 inference was executed using version 2.1.0 of the pipeline with the “monomer” parameter set, following multiple sequence alignment (MSA) construction and template structure retrieval from UniRef90 (based on UniProt 2021_04), UniClust30 (uniclust30_2018_08), MGnify (mgy_clusters_2018_12.fa), BFD (bfd_metaclust_clu_complete_id30_c90_final_seq), and PDB70 (pdb70_from_mmcif_200401) using HHblits 3.3.0 and JackHMMER from the HMMER 3.3.2 suite. For the three sequences for which the sequence retrieval pipeline in AlphaFold2 failed to complete MSA construction, ColabFold (version 1.3.0) was used to predict structures with the “AlphaFold2-ptm” weights and MMseqs2 (version 15.6f452) searches against uniref30_2103_db and colabfold_envdb_202108_db_seq. All positions corresponding to stop codons in the RT products were substituted with tryptophans (encoded by UGG).

### Prediction of IDRs with NeProc

IDRs were predicted using NeProc (Figure 4D-E) ^35^. NeProc predicted IDRs based on position-specific scoring matrices generated from PSI-BLAST searches performed with blastpgp (version 2.2.26) against the UniRef90 database (based on UniProt 2016_07). The entire polypeptides were divided into predicted IDRs and ordered regions (ORs). The percentage of IDRs was defined separately for the entire protein sequence, the ORF1-derived polypeptide, and the ORF2-derived polypeptide as the number of amino acid residues within predicted IDRs divided by the amino acid sequence length of the corresponding region.

### Gene ontology and protein feature enrichment analysis

Gene Ontology (GO) term enrichment analysis (Figure S6) and UniProtKB sequence feature enrichment analysis (Figure 4A) were performed using the Database for Annotation, Visualization, and Integrated Discovery (DAVID) ^63^. The RT gene set was used as the query list, and the background gene set was defined as both RT and non-RT genes. In this annotation category, “COMPBIAS” denotes compositionally biased protein regions, such as regions enriched in polar residues, whereas “REGION” denotes annotated protein regions, including disordered regions. Enriched terms were visualized using *R*.

### Evolutionary analysis of ORF2 and readthrough stop codons

To investigate the evolutionary origins of ORF2 sequences and RT stop codons, we analyzed homologous sequences of the 87 RT genes, whose ORF2 encodes more than 50 amino acids, across 13 arthropod species: *D. melanogaster, D. simulans, D. yakuba, D. virillis, Scaptodrosophila lebanonensis, Ceratitis capitata, Musca domestica, Megaselia abdita, Aedes aegypti, Thermobia domestica, Periplaneta americana, Chamberlinius hualienensis,* and *Centruroides sculpturatus*. The evolutionary origins of ORF2 and the RT stop codon were evaluated independently.

ORF2 conservation was assessed using both BLASTP searches against annotated proteomes and TBLASTN searches against genome assemblies. The most distantly related species in which an ORF2 homolog was detected was recorded as the oldest detectable ORF2 occurrence. RT stop codons were evaluated using protein annotations whenever possible. Because alternative splicing can produce multiple protein isoforms, all protein isoforms showing an ORF2 BLASTP hit were examined. When ORF1 and ORF2 were mapped to the same annotated protein, the corresponding ORF1–ORF2 boundary was inspected. An X at the orthologous boundary was interpreted as evidence for an RT stop codon, whereas an ordinary amino acid at this position indicated a continuous protein without an intervening stop codon.

When protein annotations were insufficient to resolve the ORF1–ORF2 boundary, genomic sequence and GFF annotations were used. The terminal stop codon of the annotated ORF1 coding sequence was identified, and the position and reading frame of downstream ORF2 TBLASTN hits were examined. RT-stop presence was considered supported when the ORF2 homolog occurred downstream of the ORF1 terminal stop in the expected readthrough frame and satisfied the genomic or transcript-based criteria used in the analysis.

Finally, to obtain direct evidence for the evolutionary conversion of a continuous protein into an RT protein, we identified genes for which the orthologous ORF1–ORF2 boundary encoded an ordinary amino acid in an evolutionarily more distant lineage but an X in a more closely related lineage. These cases provide direct protein-level evidence consistent with acquisition of an RT stop codon after the ORF2 sequence was already present.

### Construction of reporter strains and confocal imaging

DNA fragments containing *tagRFP*, ORF1/ORF2 of the target mRNAs (*Bru3-RE, CASK-RF, eEF1α2-RA, Synapsin-RE, Glut4EF-RH*), and *Venus* were cloned into the *pBFv-UAS3* plasmid (Addgene #138399). cDNA was PCR-amplified from the plasmid stocks in the *Drosophila* Genomics Resource Center (DGRC). The resultant plasmids were injected into the *y^1^v*^1^ *P{nos-phiC31};P{CaryP}attP40* embryo, screened for a *v+* phenotype, and balanced with *CyO* balancer by BestGene Inc.

These UAS strains were crossed to the pan-neuronal driver (*w*^1118^*;;R57C10-GAL4*). Brains of the F1 progeny (3-7 days old male) were manually dissected and fixed in 2% paraformaldehyde in PBS for 1hr at room temperature with gentle shaking. The brains were washed three times with PBST (0.1% Triton X-100 in PBS) and were mounted in SeeDB2S ^64^. The brains were imaged by confocal microscopy (Figures 5 and S7) (Zeiss LSM 800 and Olympus FV1200).

## References

1. Schueren, F., and Thoms, S. (2016). Functional Translational Readthrough: A Systems Biology Perspective. PLoS Genet 12, e1006196. 10.1371/journal.pgen.1006196.

2. Mangkalaphiban, K., He, F., Ganesan, R., Wu, C., Baker, R., and Jacobson, A. (2021). Transcriptome-wide investigation of stop codon readthrough in Saccharomyces cerevisiae. PLoS Genet 17, e1009538. 10.1371/journal.pgen.1009538.

3. Makarova, N., Martín de la Fuente, D., Stašuk, M., Martínez, I.O., Pöhls, J., Jungreis, I., Kellis, M., Battistini, F., Garrido Martín, D., Lescure, A., et al. (2026). Regulatory landscape of widespread stop codon readthrough in Drosophila. Preprint at bioRxiv: The Preprint Server for Biology, 10.64898/2026.06.10.731275 https://doi.org/10.64898/2026.06.10.731275.

4. Dunn, J.G., Foo, C.K., Belletier, N.G., Gavis, E.R., and Weissman, J.S. (2013). Ribosome profiling reveals pervasive and regulated stop codon readthrough in Drosophila melanogaster. eLife 2, e01179. 10.7554/eLife.01179.

5. Mangkalaphiban, K., Fu, L., Du, M., Thrasher, K., Keeling, K.M., Bedwell, D.M., and Jacobson, A. (2024). Extended stop codon context predicts nonsense codon readthrough efficiency in human cells. Nat Commun 15, 2486. 10.1038/s41467-024-46703-z.

6. Sapkota, D., Lake, A.M., Yang, W., Yang, C., Wesseling, H., Guise, A., Uncu, C., Dalal, J.S., Kraft, A.W., Lee, J.-M., et al. (2019). Cell-Type-Specific Profiling of Alternative Translation Identifies Regulated Protein Isoform Variation in the Mouse Brain. Cell Reports 26, 594–607.e7. 10.1016/j.celrep.2018.12.077.

7. Wangen, J.R., and Green, R. (2020). Stop codon context influences genome-wide stimulation of termination codon readthrough by aminoglycosides. Elife 9, e52611. 10.7554/eLife.52611.

8. Toledano, I., Supek, F., and Lehner, B. (2024). Genome-scale quantification and prediction of pathogenic stop codon readthrough by small molecules. Nat Genet 56, 1914–1924. 10.1038/s41588-024-01878-5.

9. Cridge, A.G., Crowe-McAuliffe, C., Mathew, S.F., and Tate, W.P. (2018). Eukaryotic translational termination efficiency is influenced by the 3’ nucleotides within the ribosomal mRNA channel. Nucleic Acids Res 46, 1927–1944. 10.1093/nar/gkx1315.

10. Steneberg, P., and Samakovlis, C. (2001). A novel stop codon readthrough mechanism produces functional Headcase protein in Drosophila trachea. EMBO Rep 2, 593–597. 10.1093/embo-reports/kve128.

11. Hudson, A.M., Szabo, N.L., Loughran, G., Wills, N.M., Atkins, J.F., and Cooley, L. (2021). Tissue-specific dynamic codon redefinition in *Drosophila*. Proc. Natl. Acad. Sci. U.S.A. 118, e2012793118. 10.1073/pnas.2012793118.

12. Li, C., and Zhang, J. (2019). Stop-codon read-through arises largely from molecular errors and is generally nonadaptive. PLoS Genet 15, e1008141. 10.1371/journal.pgen.1008141.

13. Müller, M.B.D., Kasturi, P., Jayaraj, G.G., and Hartl, F.U. (2023). Mechanisms of readthrough mitigation reveal principles of GCN1-mediated translational quality control. Cell 186, 3227–3244.e20. 10.1016/j.cell.2023.05.035.

14. Arribere, J.A., Cenik, E.S., Jain, N., Hess, G.T., Lee, C.H., Bassik, M.C., and Fire, A.Z. (2016). Translation readthrough mitigation. Nature 534, 719–723. 10.1038/nature18308.

15. Prieto-Godino, L.L., Rytz, R., Bargeton, B., Abuin, L., Arguello, J.R., Peraro, M.D., and Benton, R. (2016). Olfactory receptor pseudo-pseudogenes. Nature 539, 93–97. 10.1038/nature19824.

16. Karki, P., Carney, T.D., Maracci, C., Yatsenko, A.S., Shcherbata, H.R., and Rodnina, M.V. (2022). Tissue-specific regulation of translational readthrough tunes functions of the traffic jam transcription factor. Nucleic Acids Research 50, 6001–6019. 10.1093/nar/gkab1189.

17. Eswarappa, S.M., Potdar, A.A., Koch, W.J., Fan, Y., Vasu, K., Lindner, D., Willard, B., Graham, L.M., DiCorleto, P.E., and Fox, P.L. (2014). Programmed translational readthrough generates antiangiogenic VEGF-Ax. Cell 157, 1605–1618. 10.1016/j.cell.2014.04.033.

18. Zhao, Y., Lindberg, B.G., Esfahani, S.S., Tang, X., Piazza, S., and Engström, Y. (2021). Stop codon readthrough alters the activity of a POU/Oct transcription factor during Drosophila development. BMC Biol 19, 185. 10.1186/s12915-021-01106-0.

19. Sapkota, D., Florian, C., Doherty, B.M., White, K.M., Reardon, K.M., Ge, X., Garbow, J.R., Yuede, C.M., Cirrito, J.R., and Dougherty, J.D. (2022). Aqp4 stop codon readthrough facilitates amyloid-β clearance from the brain. Brain 145, 2982–2990. 10.1093/brain/awac199.

20. Mueller, S.M., McFarland White, K., Fass, S.B., Chen, S., Shi, Z., Ge, X., Engelbach, J.A., Gaines, S.H., Bice, A.R., Vasek, M.J., et al. (2023). Evaluation of gliovascular functions of AQP4 readthrough isoforms. Front Cell Neurosci 17, 1272391. 10.3389/fncel.2023.1272391.

21. Jungreis, I., Lin, M.F., Spokony, R., Chan, C.S., Negre, N., Victorsen, A., White, K.P., and Kellis, M. (2011). Evidence of abundant stop codon readthrough in *Drosophila* and other metazoa. Genome Res. 21, 2096–2113. 10.1101/gr.119974.110.

22. Jungreis, I., Chan, C.S., Waterhouse, R.M., Fields, G., Lin, M.F., and Kellis, M. (2016). Evolutionary Dynamics of Abundant Stop Codon Readthrough. Mol Biol Evol 33, 3108–3132. 10.1093/molbev/msw189.

23. Ichinose, T., Kanno, M., Makino, T., Shichino, Y., Iwasaki, S., and Tanimoto, H. (2026). Activity-dependent synthesis of translational machinery through dynamic remodeling of ribonucleoprotein granules. Preprint at Neuroscience, 10.64898/2026.06.28.734671 https://doi.org/10.64898/2026.06.28.734671.

24. Ichinose, T., Kondo, S., Kanno, M., Shichino, Y., Mito, M., Iwasaki, S., and Tanimoto, H. (2024). Translational regulation enhances distinction of cell types in the nervous system. eLife 12, RP90713. 10.7554/eLife.90713.

25. Ichinose, T., and Tanimoto, H. (2025). Profiling translation in the nervous system. J Biochem 177, 239–246. 10.1093/jb/mvae096.

26. Chen, X., and Dickman, D. (2017). Development of a tissue-specific ribosome profiling approach in Drosophila enables genome-wide evaluation of translational adaptations. PLoS Genet 13, e1007117. 10.1371/journal.pgen.1007117.

27. Simonyan, K., Vedaldi, A., and Zisserman, A. (2014). Deep Inside Convolutional Networks: Visualising Image Classification Models and Saliency Maps. Preprint at arXiv, 10.48550/arXiv.1312.6034 https://doi.org/10.48550/arXiv.1312.6034.

28. Baehrens, D., Schroeter, T., Harmeling, S., Kawanabe, M., Hansen, K., and Müller, K.-R. (2010). How to Explain Individual Classification Decisions. Journal of Machine Learning Research 11.

29. Gruber, A.R., Lorenz, R., Bernhart, S.H., Neuböck, R., and Hofacker, I.L. (2008). The Vienna RNA websuite. Nucleic Acids Res 36, W70–74. 10.1093/nar/gkn188.

30. Lorenz, R., Bernhart, S.H., Höner Zu Siederdissen, C., Tafer, H., Flamm, C., Stadler, P.F., and Hofacker, I.L. (2011). ViennaRNA Package 2.0. Algorithms Mol Biol 6, 26. 10.1186/1748-7188-6-26.

31. Schuller, A.P., and Green, R. (2018). Roadblocks and resolutions in eukaryotic translation. Nat Rev Mol Cell Biol 19, 526–541. 10.1038/s41580-018-0011-4.

32. Ingolia, N.T., Ghaemmaghami, S., Newman, J.R.S., and Weissman, J.S. (2009). Genome-wide analysis in vivo of translation with nucleotide resolution using ribosome profiling. Science 324, 218–223. 10.1126/science.1168978.

33. Jumper, J., Evans, R., Pritzel, A., Green, T., Figurnov, M., Ronneberger, O., Tunyasuvunakool, K., Bates, R., Žídek, A., Potapenko, A., et al. (2021). Highly accurate protein structure prediction with AlphaFold. Nature 596, 583–589. 10.1038/s41586-021-03819-2.

34. Mirdita, M., Schütze, K., Moriwaki, Y., Heo, L., Ovchinnikov, S., and Steinegger, M. (2022). ColabFold: making protein folding accessible to all. Nat Methods 19, 679–682. 10.1038/s41592-022-01488-1.

35. Anbo, H., Amagai, H., and Fukuchi, S. (2020). NeProc predicts binding segments in intrinsically disordered regions without learning binding region sequences. Biophys Physicobiol 17, 147–154. 10.2142/biophysico.BSJ-2020026.

36. Wilson, B.A., Foy, S.G., Neme, R., and Masel, J. (2017). Young Genes are Highly Disordered as Predicted by the Preadaptation Hypothesis of De Novo Gene Birth. Nat Ecol Evol 1, 0146–0146. 10.1038/s41559-017-0146.

37. Banerjee, S., and Chakraborty, S. (2017). Protein intrinsic disorder negatively associates with gene age in different eukaryotic lineages. Mol Biosyst 13, 2044–2055. 10.1039/c7mb00230k.

38. Peng, J., and Zhao, L. (2024). The origin and structural evolution of de novo genes in Drosophila. Nat Commun 15, 810. 10.1038/s41467-024-45028-1.

39. Van Roey, K., Uyar, B., Weatheritt, R.J., Dinkel, H., Seiler, M., Budd, A., Gibson, T.J., and Davey, N.E. (2014). Short linear motifs: ubiquitous and functionally diverse protein interaction modules directing cell regulation. Chem Rev 114, 6733–6778. 10.1021/cr400585q.

40. Tompa, P., Davey, N.E., Gibson, T.J., and Babu, M.M. (2014). A million peptide motifs for the molecular biologist. Mol Cell 55, 161–169. 10.1016/j.molcel.2014.05.032.

41. Kansara, L., Wolfstetter, G., Wintermayr, D., Alexopoulos, I., Escós, A., Friedländer, M.R., and Engström, Y. (2025). The mRNA architecture of the termination site primes programmed stop codon readthrough events in *Drosophila*. Preprint at Molecular Biology, 10.1101/2025.11.07.685291 https://doi.org/10.1101/2025.11.07.685291.

42. Han, P., Shichino, Y., Schneider-Poetsch, T., Mito, M., Hashimoto, S., Udagawa, T., Kohno, K., Yoshida, M., Mishima, Y., Inada, T., et al. (2020). Genome-wide Survey of Ribosome Collision. Cell Reports 31, 107610. 10.1016/j.celrep.2020.107610.

43. Meydan, S., and Guydosh, N.R. (2020). Disome and Trisome Profiling Reveal Genome-wide Targets of Ribosome Quality Control. Mol Cell 79, 588–602.e6. 10.1016/j.molcel.2020.06.010.

44. Zhao, T., Chen, Y.-M., Li, Y., Wang, J., Chen, S., Gao, N., and Qian, W. (2021). Disome-seq reveals widespread ribosome collisions that promote cotranslational protein folding. Genome Biol 22, 16. 10.1186/s13059-020-02256-0.

45. Roy, B., Leszyk, J.D., Mangus, D.A., and Jacobson, A. (2015). Nonsense suppression by near-cognate tRNAs employs alternative base pairing at codon positions 1 and 3. Proc Natl Acad Sci U S A 112, 3038–3043. 10.1073/pnas.1424127112.

46. Carvunis, A.-R., Rolland, T., Wapinski, I., Calderwood, M.A., Yildirim, M.A., Simonis, N., Charloteaux, B., Hidalgo, C.A., Barbette, J., Santhanam, B., et al. (2012). Proto-genes and de novo gene birth. Nature 487, 370–374. 10.1038/nature11184.

47. Kosinski, L.J., and Masel, J. (2020). Readthrough Errors Purge Deleterious Cryptic Sequences, Facilitating the Birth of Coding Sequences. Molecular Biology and Evolution 37, 1761–1774. 10.1093/molbev/msaa046.

48. Uversky, V.N. (2013). The most important thing is the tail: multitudinous functionalities of intrinsically disordered protein termini. FEBS Lett 587, 1891–1901. 10.1016/j.febslet.2013.04.042.

49. Romero, P.R., Zaidi, S., Fang, Y.Y., Uversky, V.N., Radivojac, P., Oldfield, C.J., Cortese, M.S., Sickmeier, M., LeGall, T., Obradovic, Z., et al. (2006). Alternative splicing in concert with protein intrinsic disorder enables increased functional diversity in multicellular organisms. Proc Natl Acad Sci U S A 103, 8390–8395. 10.1073/pnas.0507916103.

50. Kumar, M., Michael, S., Alvarado-Valverde, J., Zeke, A., Lazar, T., Glavina, J., Nagy-Kanta, E., Donagh, J.M., Kalman, Z.E., Pascarelli, S., et al. (2024). ELM—the Eukaryotic Linear Motif resource—2024 update. Nucleic Acids Research 52, D442–D455. 10.1093/nar/gkad1058.

51. Schad, E., Fichó, E., Pancsa, R., Simon, I., Dosztányi, Z., and Mészáros, B. (2018). DIBS: a repository of disordered binding sites mediating interactions with ordered proteins. Bioinformatics 34, 535–537. 10.1093/bioinformatics/btx640.

52. Fukuchi, S., Amemiya, T., Sakamoto, S., Nobe, Y., Hosoda, K., Kado, Y., Murakami, S.D., Koike, R., Hiroaki, H., and Ota, M. (2014). IDEAL in 2014 illustrates interaction networks composed of intrinsically disordered proteins and their binding partners. Nucl. Acids Res. 42, D320–D325. 10.1093/nar/gkt1010.

53. Picchio, L., Legagneux, V., Deschamps, S., Renaud, Y., Chauveau, S., Paillard, L., and Jagla, K. (2018). Bruno-3 regulates sarcomere component expression and contributes to muscle phenotypes of myotonic dystrophy type 1. Dis Model Mech 11, dmm031849. 10.1242/dmm.031849.

54. Delaunay, J., Le Mée, G., Ezzeddine, N., Labesse, G., Terzian, C., Capri, M., and Aït-Ahmed, O. (2004). The Drosophila Bruno paralogue Bru-3 specifically binds the EDEN translational repression element. Nucleic Acids Res 32, 3070–3082. 10.1093/nar/gkh627.

55. Dasgupta, T., and Ladd, A.N. (2012). The importance of CELF control: molecular and biological roles of the CUG-BP, Elav-like family of RNA-binding proteins. Wiley Interdiscip Rev RNA 3, 104–121. 10.1002/wrna.107.

56. Oas, S.T., Bryantsev, A.L., and Cripps, R.M. (2014). Arrest is a regulator of fiber-specific alternative splicing in the indirect flight muscles of Drosophila. J Cell Biol 206, 895–908. 10.1083/jcb.201405058.

57. Suzuki, H., Jin, Y., Otani, H., Yasuda, K., and Inoue, K. (2002). Regulation of alternative splicing of alpha-actinin transcript by Bruno-like proteins. Genes Cells 7, 133–141. 10.1046/j.1356-9597.2001.00506.x.

58. Rashad, S., Kitamura, Y., Nagai, T., Ando, D., Mousa, A., Ikenouchi, H., Endo, H., and Niizuma, K. (2025). Ischemic stroke induces staged translational remodeling in mouse cortex independent from transcriptional responses. Preprint at Molecular Biology, 10.1101/2025.08.15.670488 https://doi.org/10.1101/2025.08.15.670488.

59. Sun, H., Nishioka, T., Hiramatsu, S., Kondo, S., Amano, M., Kaibuchi, K., Ichinose, T., and Tanimoto, H. (2020). Dopamine Receptor Dop1R2 Stabilizes Appetitive Olfactory Memory through the Raf/MAPK Pathway in Drosophila. J. Neurosci. 40, 2935–2942. 10.1523/JNEUROSCI.1572-19.2020.

60. Alipanahi, B., Delong, A., Weirauch, M.T., and Frey, B.J. (2015). Predicting the sequence specificities of DNA- and RNA-binding proteins by deep learning. Nat Biotechnol 33, 831–838. 10.1038/nbt.3300.

61. Kelley, D.R., Snoek, J., and Rinn, J.L. (2016). Basset: learning the regulatory code of the accessible genome with deep convolutional neural networks. Genome Res 26, 990–999. 10.1101/gr.200535.115.

62. Kingma, D.P., and Ba, J. (2017). Adam: A Method for Stochastic Optimization. Preprint at arXiv, 10.48550/arXiv.1412.6980 https://doi.org/10.48550/arXiv.1412.6980.

63. Dennis, G., Sherman, B.T., Hosack, D.A., Yang, J., Gao, W., Lane, H.C., and Lempicki, R.A. (2003). DAVID: Database for Annotation, Visualization, and Integrated Discovery. Genome Biol 4, R60. 10.1186/gb-2003-4-9-r60.

64. Ke, M.-T., Nakai, Y., Fujimoto, S., Takayama, R., Yoshida, S., Kitajima, T.S., Sato, M., and Imai, T. (2016). Super-Resolution Mapping of Neuronal Circuitry With an Index-Optimized Clearing Agent. Cell Rep 14, 2718–2732. 10.1016/j.celrep.2016.02.057.

65. Ioffe, S., and Szegedy, C. (2015). Batch Normalization: Accelerating Deep Network Training by Reducing Internal Covariate Shift. Preprint at arXiv, 10.48550/arXiv.1502.03167 https://doi.org/10.48550/arXiv.1502.03167.

66. Srivastava, N., Hinton, G., Krizhevsky, A., Sutskever, I., and Salakhutdinov, R. (2014). Dropout: A Simple Way to Prevent Neural Networks from Overfitting. Journal of Machine Learning Research 15.

