## Supplementary figures for "Neuronal stop-codon readthrough is associated with ribosome pausing and appends lineage-restricted extensions that can alter protein localization in *Drosophila*"

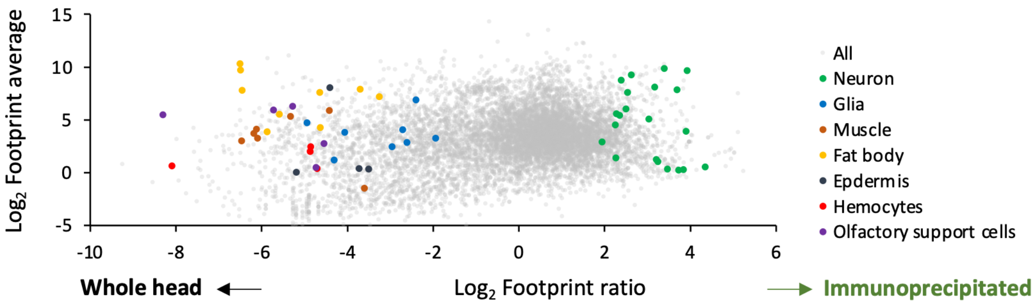


**Figure S1. Neuron-specific ribosome purification enriches neuronal markers.**

Scatter plot comparing ribosome-footprint abundance and enrichment between whole-head and immunoprecipitated neuron-specific ribosome profiling samples. Each point represents one gene. The x-axis shows the $\log_2$(footprint ratio) between immunoprecipitated and whole-head samples, and the y-axis shows the average $\log_2$(footprint abundance). Marker genes are as follows.

Neuron: *nSyb, Ilp2, Trh, Shaw, nAChRa1, FoxP, Gad1, ChAT, Vglut, para, Sh, Tbh, ppk, AstC, rad, NPF, mAChR-B, elav, Rab3, Syta, Hug*.

Glia: *alrm, repo, moody, wrapper, nrv2, zyd, Gat, Balat*.

Muscle: *Mhc, wupA, Tm2, sls, up, Mp20, Mlp60A*.

Fat body: *Lsd-2, FASN1, Yp1, Yp2, apolpp, Lsp2, fit, AkhR, fon*.

Epidermis: *Cpr47Ea, Cpr49Af, Cpr65Au, Cpr67Fb*.

Hemocytes: *Hml, eater, NimC1, NimC2*.

Olfactory support cells: *Obp19b, Obp49a, Obp56g, Obp83ef, Obp99a*.

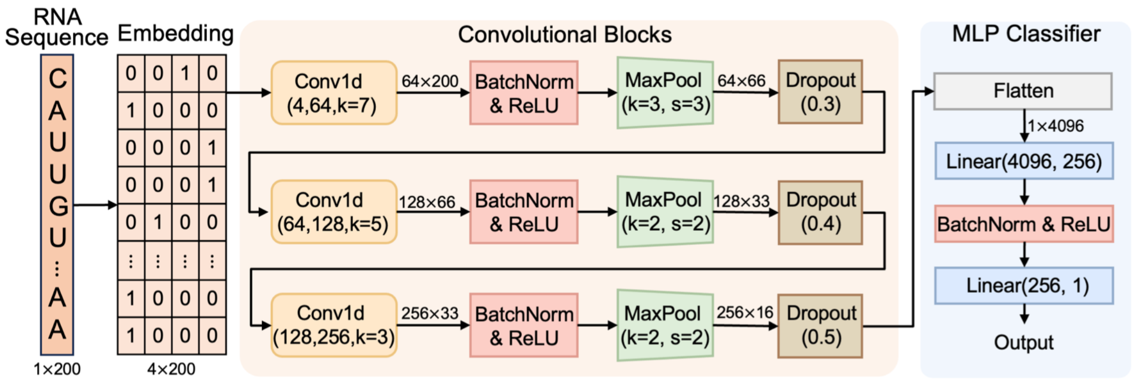


**Figure S2. Architecture of the 1D convolutional neural network.**

Detailed architecture of the sequence-based CNN classifier. Input RNA sequences spanning 200 nt around the annotated stop codon are converted to binary matrices and passed through three convolutional blocks. The first block used Conv1D with 64 filters and kernel size 7, followed by batch normalization, ReLU activation, max pooling, and dropout [65,66]. The second and third blocks used 128 and 256 filters with kernel sizes 5 and 3, respectively, followed by batch normalization, ReLU activation, max pooling, and dropout. The resulting feature vector was flattened and passed through a multilayer perceptron classifier to predict RT versus non-RT status. Dropout probabilities were 0.3, 0.4, and 0.5 for the three convolutional blocks.

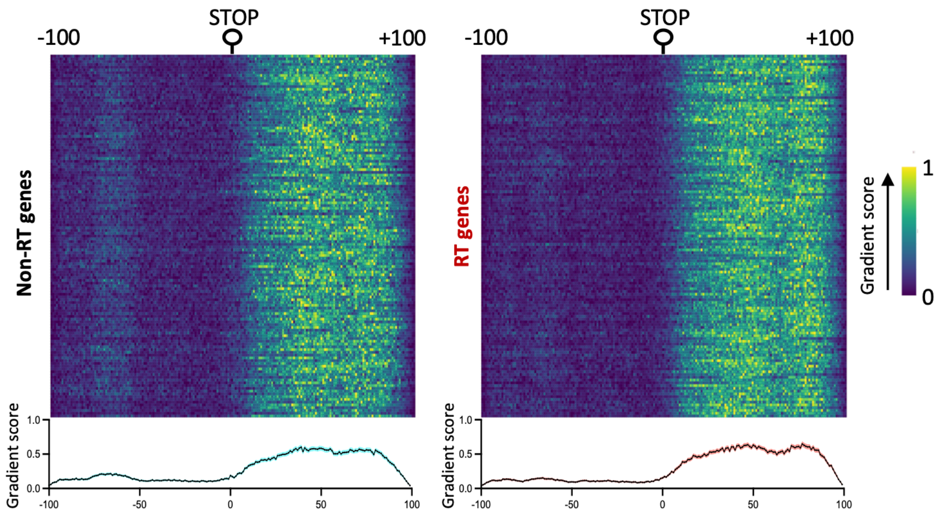


**Figure S3. CNN attribution analysis on the training transcript set.**

Gradient-based attribution analysis of the CNN classifier applied to the training set. Heatmaps show nucleotide-level gradient scores for non-RT and RT transcripts. Line plots at the bottom show the average ± standard error across positions.

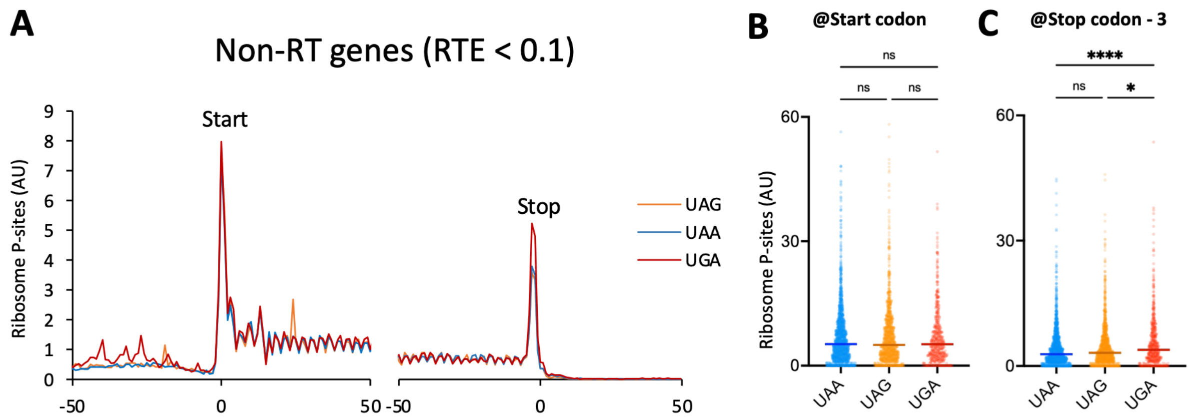


**Figure S4. UGA induces modest stop-codon pausing even in non-RT transcripts.**

(A) Metagene analysis of ribosome P-sites around start and stop codons in non-RT transcripts, stratified by stop-codon identity.

(B, C) Quantification of ribosome P-site density at the start codon (B) and at the position 3 nt upstream of the annotated stop codon (C), corresponding to a ribosome with the stop codon positioned in the A-site.

Dunn's multiple comparison test is used, and center lines indicate the median. ns, not significant; *$P < 0.05$; ****$P < 0.0001$.

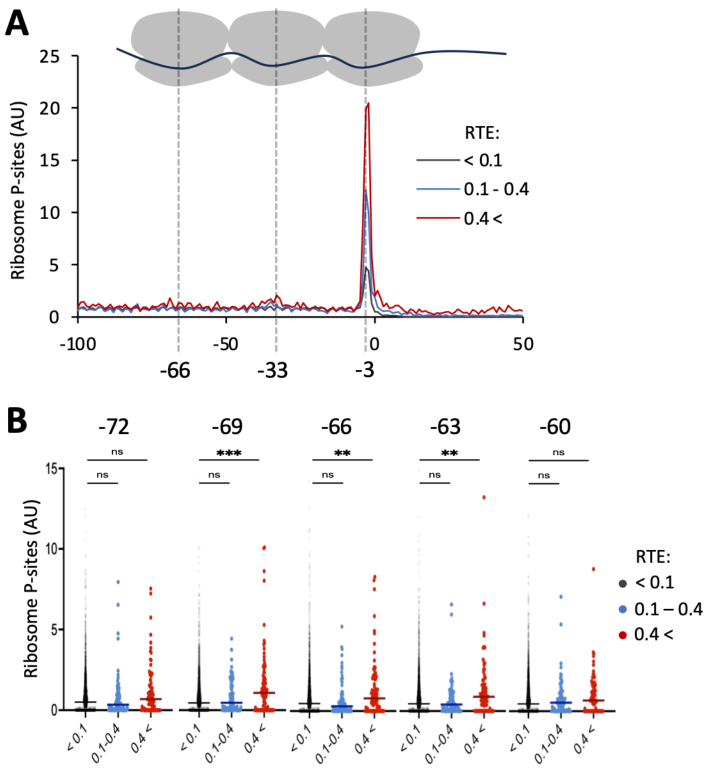


**Figure S5. Upstream ribosome-footprint peaks support ribosome queuing on RT transcripts.**

(A) Metagene analysis of ribosome P-sites upstream of annotated stop codons, stratified by RTE. In addition to the major peak near the stop codon, RT transcripts show secondary and tertiary upstream peaks near -39 and −69 nt.

(B) Quantification of ribosome P-site density at −72, −69, −66, −63, and −60 nt.

Dunn's multiple comparison test is used, and center lines indicate the median. ns, not significant; $^{**}P < 0.01$; $^{***}P < 0.001$.

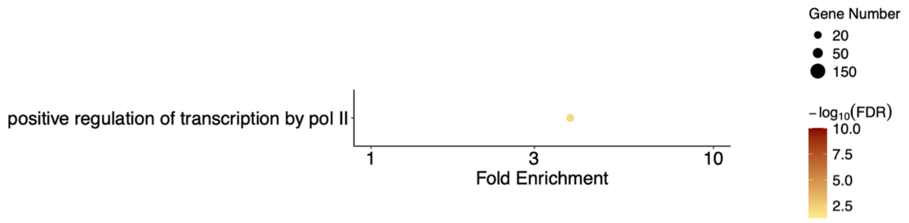


**Figure S6. Functional enrichment analysis of RT genes.**

Gene Ontology enrichment analyses of RT genes. GO analysis showed limited enrichment of molecular or biological functions, with enrichment detected for “positive regulation of transcription by pol II” (GO:0045944).

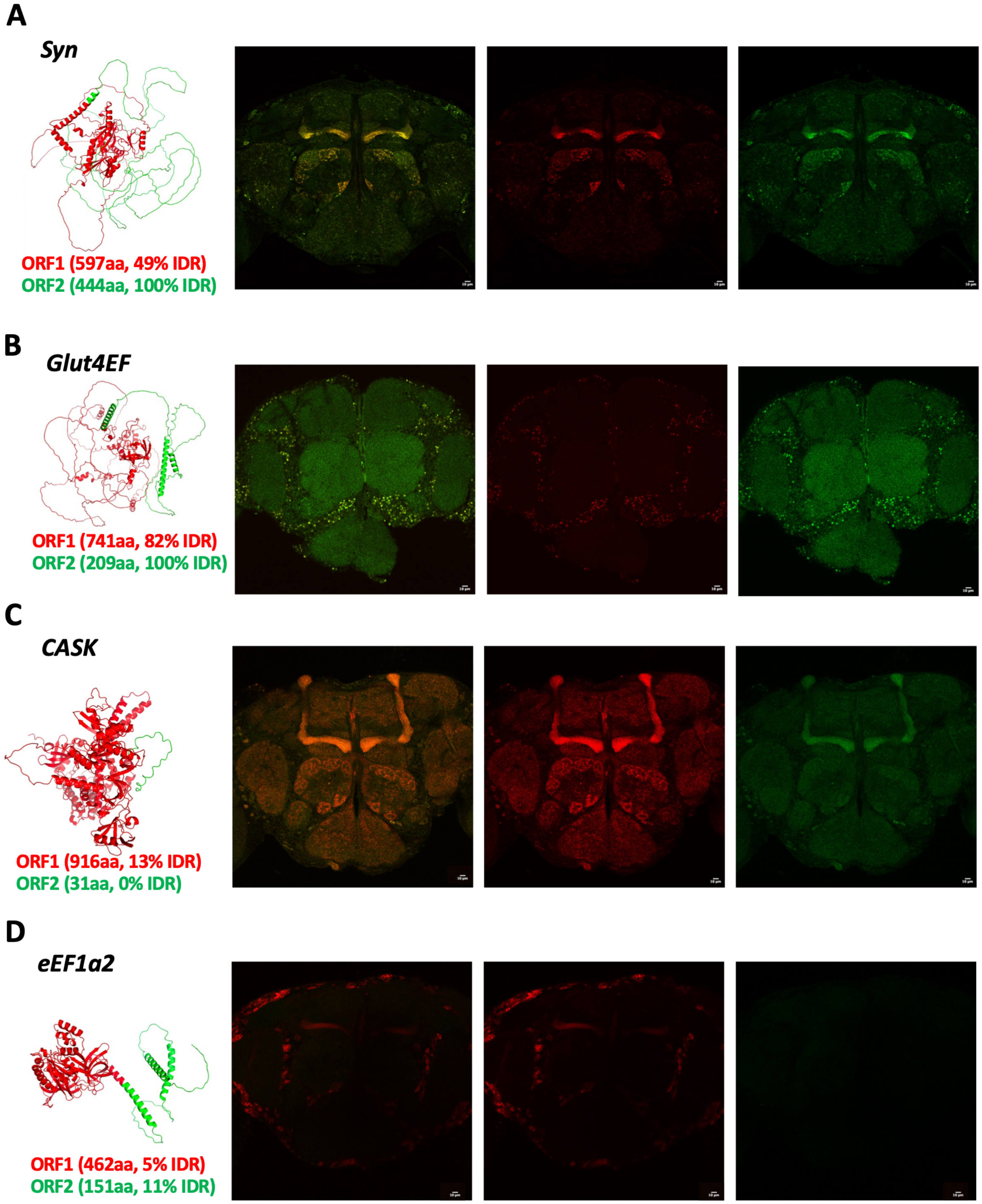


**Figure S7. Dual-color reporters for other genes.**

Representative confocal whole-brain images of dual-color readthrough reporters for *Synapsin-RE, Glut4EF-RH, CASK-RF,* and *eEF1α2-RA*. Predicted protein structure (AlphaFold2) is shown on the left (green: ORF1, red: ORF2). Polypeptide length of ORF1 and ORF2, and predicted IDR fraction (NeProc) are noted. Reporters were expressed in neurons using *nSyb-GAL4*. Scale bars, 10 μm.
